# Targeting SEZ6L2 in Colon Cancer: Efficacy of Bexarotene and Implications for Survival

**DOI:** 10.1101/2024.06.19.599786

**Authors:** Huajun Zheng, Jianying Zheng, Yan Shen

## Abstract

**Background:** Bexarotene, also recognized as Targretin, is categorized as a retinoid, a type of cancer drug. Nevertheless, the precise mechanisms of Bexarotene in relation to colon cancer remain unclear. In colon cancer, SEZ6L2 was suggested as one of the biomarkers and targets. This study presents a comprehensive exploration of the role of SEZ6L2 in colon cancer.

**Methods:** We utilized both TCGA data and a cohort of Chinese patients. In a meticulous analysis of 478 colon cancer cases, SEZ6L2 expression levels were examined in relation to clinical characteristics, staging parameters, and treatment outcomes. Additionally, we investigated the pharmacological impact of Bexarotene on SEZ6L2, demonstrating a significant downregulation of SEZ6L2 at both mRNA and protein levels in colon cancer patients following Bexarotene treatment.

**Results:** SEZ6L2 consistently overexpresses in colon cancer, serving as a potential universal biomarker with prognostic significance, validated in a diverse Chinese cohort. In vitro, SEZ6L2 promotes cell viability without affecting migration. Bexarotene treatment inhibits SEZ6L2 expression, correlating with reduced viability both in vitro and in vivo. SEZ6L2 overexpression accelerates declining survival rates in an in vivo context. Bexarotene’s efficacy is context-dependent, effective in parental cells but not with SEZ6L2 overexpression. Computational predictions suggest a direct SEZ6L2-Bexarotene interaction, warranting further experimental exploration.

**Conclusion:** The study provides valuable insights into SEZ6L2 as a prognostic biomarker in colon cancer, revealing its intricate relationship with clinical parameters, treatment outcomes, and Bexarotene effects. Context-dependent therapeutic responses emphasize the nuanced understanding required for SEZ6L2’s role in colon cancer, paving the way for targeted therapeutic strategies.

## 1. Introduction

Colon cancer, a complex and heterogeneous disease, continues to pose significant challenges in both diagnosis and treatment. It stands as the third most prevalent cancer globally and is the second leading cause of cancer-related deaths[1]. Distant metastasis, particularly in the liver, is the primary cause of fatality in colon cancer patients. At the time of initial colon cancer diagnosis, approximately 25% of patients already exhibit distant metastatic disease, and more than 50% eventually develop metastases[2]. The metastatic process in colorectal cancer is intricate, involving multiple biological stages such as tumor cell shedding from the primary site, intravasation, survival of circulating tumor cells, extravasation, and eventual colonization at a distant site[3]. Recent clinical studies have highlighted the association between mutations in key driver genes like KRAS, BRAF, and p53 and metastatic colorectal cancer[4]. The identification of novel biomarkers and therapeutic targets is paramount for advancing our understanding of this malignancy and improving patient outcomes.

SEZ6L2 is a protein associated with seizures and is situated on the cell surface. Positioned in the chromosome 16p11.2 region, which is considered to harbor candidate genes linked to autism spectrum disorders (ASD), there is currently no direct evidence connecting this gene to ASD. However, it is noteworthy that elevated expression of SEZ6L2 has been observed in cancer such as lung cancer[5] and breast cancer [6], designating the protein as a new prognostic marker for lung cancer. Additionally, alternative splicing of this gene gives rise to various transcript variants. In colon cancer, SEZ6L2 was suggested as one of the transcriptome signature for cancer prognostic[7]. This study focuses on the multifaceted role of SEZ6L2, a cell adhesion molecule, in the context of colon cancer, integrating comprehensive analyses from clinical databases, in vitro assays, and in vivo models.

Bexarotene, also recognized as Targretin, is categorized as a retinoid, a type of cancer drug. It is employed in the treatment of advanced skin lymphomas, specifically cutaneous T cell lymphomas, which encompass mycosis fungoides and Sezary syndrome. The pronunciation of bexarotene is commonly rendered as “becks-a-roh-teen.” Regarding its mechanism of action, bexarotene operates as a retinoid, belonging to the category of drugs associated with vitamin A[8,9]. Its mode of function involves impeding or halting the excessive growth of normal cells. Bexarotene has been used to treat colon cancer in previous studies[10,11]. Nevertheless, the precise mechanisms of Bexarotene in relation to colon cancer remain unclear. Specifically, it is not well-established whether Bexarotene has a direct impact on colon cancer cells.

Our investigation commences with an exploration of SEZ6L2’s clinical relevance, drawing on analyses from large-scale databases. We demonstrate a consistent and robust overexpression of SEZ6L2 in colon cancer tissues compared to normal counterparts, establishing its potential as a diagnostic biomarker. A comprehensive ROC analysis underscores its diagnostic accuracy, while survival analysis implicates SEZ6L2 as a promising prognostic indicator, correlating higher expression with a significantly lower survival rate. Despite the lack of associations with conventional clinical parameters, SEZ6L2 emerges as a pivotal player in cancer development and progression. Our study unveils the multifaceted role of SEZ6L2 in colon cancer, positioning it as a promising diagnostic, prognostic, and potentially therapeutic target. The comprehensive approach, encompassing clinical data, pharmacological interventions, and molecular predictions, contributes to a deeper understanding of SEZ6L2’s involvement in colon cancer, paving the way for targeted interventions and personalized treatment strategies in the clinical landscape.

## 2. Methods

### 2.1. Bioinformatics Analysis

This study utilized the TCGA COAT cohort to analyze colorectal cancer patients. Furthermore, our independent cohort consisted of colon cancer patients with surgically excised colon cancer (n=77) and adjacent non-tumor tissues (n=49) from the The Second Affiliated Hospital of Zhejiang University of Traditional Chinese Medicine. Among the patients with cancer samples collected, 50 had recorded survival data, and thus, they were subjected to survival analysis. Additionally, we included a third cohort of patients with samples collected both before and after Bexarotene treatment (>90 days). All our samples were collected following freezing, and SEZ6L2 expression was assessed using QPCR and Western blotting. The study was conducted in accordance with the Declaration of Helsinki of the World Medical Association and obtained approval from the human ethics committee of the The Second Affiliated Hospital of Zhejiang University of Traditional Chinese Medicine.

### 2.2. Cell Cultures

The HCT-116 and HCT15 cell lines were obtained from the Cell Bank of Shanghai Institute of Biochemistry and Cell Biology, Chinese Academy of Sciences (Shanghai, China). Cells were cultured in DMEM/HIGH GLUCOSE (Cytiva) supplemented with 10% FBS (LONSA) at 37 °C with 5% CO2.

### 2.3. Cellular Transfection

Cell transfection procedures were conducted following established methodologies as described in prior studies[12–15]. The overexpression plasmid were obtained from Genepharma (China). The transfection process employed Lipofectamine 3000 reagent (Invitrogen, USA) and lasted for 48 hours. The sable cell line expressing SEZ6L2 were obtain by transfection following antibiotic selection. Cells were transfected with a construct incorporating SEZ6L2 and a G418 resistance gene. Following transfection, cells were subjected to G418 selection to generate a stable cell line expressing SEZ6L2.

### 2.4. Cell Viability and Proliferation Assay

The CCK-8 (TargetMol, USA) assay assessed cell viability. HCT-116 cells were seeded into a 96-well culture plate (Thermo) at 50% confluency for 24 h. Cells were then treated with a vehicle or serial dilutions of 31 for 48 h. CCK-8 solution was added, and absorbance was measured at 450 nm by a microplate reader (Molecular Devices, CA, USA). For the colony formation assay, cells were seeded into 6-well plates (Corning) at a density of 1 × 103/well, treated with various concentrations of 31 for 48 h, and refreshed with growth medium every other day. After two weeks, cells were fixed, stained with crystal violet solution (Sigma-Aldrich), and colony images were manually captured.

### 2.5. Cell Migration Assay

Cells (1.2 × 10^4^ cells/24-well plate) were cultured in FBS-free medium and treated with drugs. Migration was monitored using the scratch wound healing assay after 24 h incubation. Images were captured using an Evos XL Core Imaging System microscope (Thermo Fisher, USA), and migration was calculated using Image J software.

### 2.6. Quantitative Reverse Transcription PCR (RT-qPCR)

Cells or tissues samples were ground in liquid nitrogen. Total RNA was extracted using RNAiso plus (Takara, Japan). RNA (1 μg) was reverse transcribed to cDNA using reverse transcriptase (Vazyme, China). Real-time PCR was performed in StepOnePlus™ Real-Time PCR Systems (Applied Biosystems, USA). The PCR conditions were 5 min at 95 °C, followed by 40 cycles of 95 °C for 10 s and 65 °C for 30 s. The mRNA expression levels were normalized with GAPDH. The primers used are listed below: SEZ6L2 forward primer, 5’-ATGAAGCTGGGGATACGC-3’; SEZ6L2 reverse primer, 5’-CCTCGTGGGATAGGGAGA-3’; GAPDH forward primer, 5’-GGACCTGACCTGCCGTCTAG-3’; and GAPDH reverse primer, 5’-GTAGCCCAGGATGC CCTT-3’.

### 2.7. Western Blotting Assay

Cells or tissues samples were harvested, washed with PBS, and total proteins were extracted with 1 × RIPA lysis buffer. Protein concentration was determined by the BCA method. Equivalent samples of cell lysates were separated by SDS-PAGE, transferred to the PVDF membrane, and incubated with specific antibodies. Detection was performed with secondary isotype-specific antibodies tagged with horseradish peroxidase, and immunocomplexes were visualized using enhanced chemiluminescence. Antibodies used include Recombinant Human SEZ6L2 protein (ab161979) and Anti-GAPDH antibody [6C5] - Loading Control (ab8245).

### 2.8. Antitumor Activity In Vivo Experiment

The animal model were developed as a previous study[16]. Female BALB/c nude mice, aged six weeks, were subcutaneously injected with either HCT-116 cells or the stable HCT-116 cell line. Our experimental design included four distinct sets of animal experiments:

1. Comparison of HCT-116 Cells and Stable HCT-116 Cell Line Expressing SEZ6L2:

- Mice received injections of HCT-116 cells and the stable HCT-116 cell line expressing SEZ6L2.
- Survival rates were recorded for both groups.
2. Bexarotene Treatment:

- Mice injected with HCT-116 cells were treated with Bexarotene.
- Survival rates were monitored.
3. Combined Treatment with HCT-116 Cells and Bexarotene:

- Mice received injections of HCT-116 cells and were subsequently treated with Bexarotene.
- After a 10-week treatment period, mice were sacrificed.
- Tumor samples were collected for pathological sections, quantitative polymerase chain reaction (QPCR), and immunohistochemical analysis.
4. Drug Treatment Groups:

- When tumor volume reached 100–150 mm3, mice were divided into four groups:

- Control: Bexarotene administered orally at a dose of 0 mg/kg/day.
- Low Dose (Low): Bexarotene administered orally at a dose of 20 mg/kg/day.
- Medium Dose (Medium): Bexarotene administered orally at a dose of 50 mg/kg/day.
- High Dose (High): Bexarotene administered orally at a dose of 100 mg/kg/day.
- Capecitabine was used as a positive control, administered orally at a dose of 100 mg/kg/day.
- Survival rates were recorded, Tumor samples were collected for pathological sections, quantitative polymerase chain reaction (QPCR), and immunohistochemical analysis.

### 2.9. Statistical Analysis

Data were presented as mean ± standard deviation (SD). Both bioinformatics and experimental data, Student’s t-test and one-way ANOVA test were applied to analyze the significance of differences. The bioinformatics data were analyzed using R software with “statistic” pack. For experiments, data were analyzed using GraphPad Prism Software 8. *P < 0.05, **P < 0.01, and ***P < 0.001 were considered statistically significant.

## 3. Results

### 3.1. SEZ6L2 is a potential target for colon cancer

This study delves into the role of SEZ6L2 in colon cancer through an analysis of open databases, including TCGA and the GTEx database. TCGA cohort information was provided in supplementary materials. The investigation reveals a significant overexpression of SEZ6L2 in colon cancer tissues compared to normal colon tissues. This finding is robustly confirmed through paired comparisons of cancer and non-cancer tissues from the same patients, establishing SEZ6L2 as a prominent colon cancer biomarker.

To assess the diagnostic potential of SEZ6L2, a comprehensive ROC analysis was conducted, yielding an AUC exceeding 80. This high diagnostic accuracy underscores the viability of SEZ6L2 as a diagnostic biomarker for colon cancer, with implications for clinical applications.

Furthermore, survival analysis indicates a noteworthy correlation between high SEZ6L2 expression and a significantly lower survival rate in patients with colon cancer. This association positions SEZ6L2 as a promising prognostic biomarker. The dual diagnostic and prognostic capabilities of SEZ6L2 underscore its potential clinical utility.

Given its selective overexpression in cancerous tissues and its correlation with patient prognosis, we propose that SEZ6L2 may serve as a specific therapeutic target for colon cancer. The observed association with cancer development, invasiveness, and progression further accentuates the research value of SEZ6L2 in the context of colon cancer.

Our study highlights SEZ6L2 as a compelling biomarker for colon cance diagnosis and prognosis, opening avenues for its potential application in clinical settings and as a target for therapeutic interventions. The multifaceted role of SEZ6L2 in colon cancer underscores its significance and warrants further exploration to enhance our understanding of its mechanistic involvement in the disease. Further analysis across different Consensus Molecular Subtypes (CMS) revealed that the survival impact of SEZ6L2 is observed in all four subtypes, with CMS3 showing the most remarkable difference.(Figure 1)

**Figure 1.**
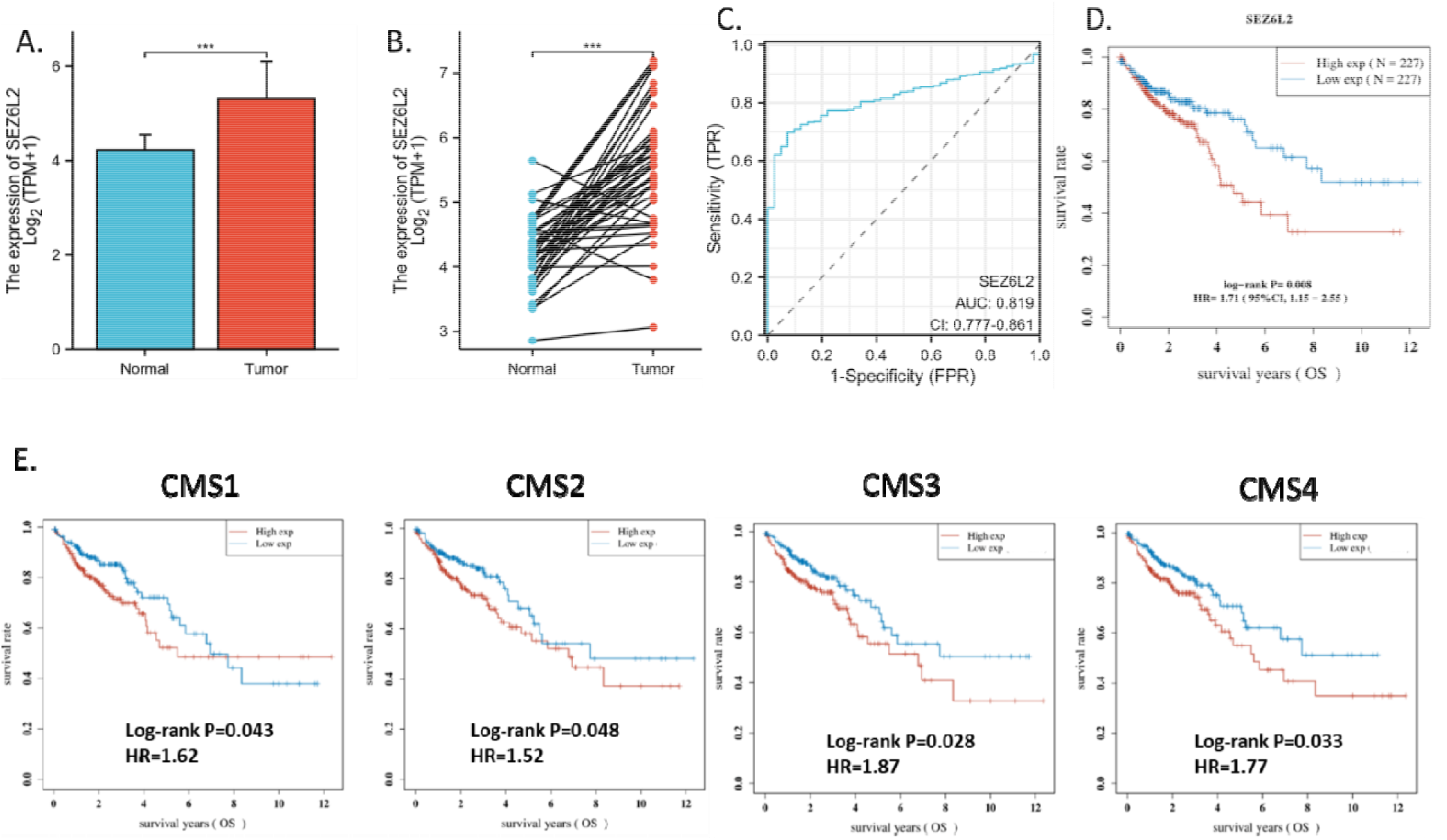
Clinical relevance of SEZ6L2 for colon cancer. A. Expression of SEZ6L2 in cancer (n=478) and non-cancer (n=249) colon tissues. (TCGA-GTEx non-paired comparison). B. Expression of SEZ6L2 in cancer and non-cancer colon tissues (n=41). (TCGA paired comparison). ***p<0.001 C. Diagnostic ROC analysis of SEZ6L2 in colon cancer (TCGA-GTEx non-paired samples). D. Survival analysis of SEZ6L2 in colon cancer(TCGA COAD samples, n=454). E. Survival analysis of SEZ6L2 in colon cancer Consensus Molecular subtypes (TCGA COAD samples).

### 3.2. Validation of the survival association of SEZ6L2 in colon cancer

Our investigation into SEZ6L2 expression in colon cancer was not only substantiated by the TCGA data but also further validated through the analysis of a cohort of Chinese patients from our hospital. Noteworthy differences were identified, emphasizing the importance of considering ethnic diversity and employing multi-dimensional analyses.

In our cohort of Chinese patients, consisting of 77 colon cancer and 49 normal colon tissues, SEZ6L2 exhibited a significant overexpression in cancerous tissues compared to normal counterparts. At the mRNA level, the average fold change was approximately 1.6. Importantly, large-scale Western blotting assays revealed a more substantial overexpression of SEZ6L2 at the protein level, with a fold change of approximately 4 in colon cancer compared to normal colon tissue. This discrepancy between mRNA and protein levels highlights the potential functional significance of SEZ6L2 at the protein level in the context of colon cancer. The inclusion of our Chinese patient cohort addressed a limitation inherent in TCGA data, predominantly derived from white patients. Our findings underscore the importance of considering ethnic diversity in cancer research, as molecular profiles may exhibit variations across different populations.

For survival analysis, consistent with the TCGA findings, survival analysis within our Chinese patient cohort revealed a compelling association between SEZ6L2 expression levels and patient outcomes. Patients with higher SEZ6L2 expression demonstrated a significantly lower survival rate compared to those with lower expression, further affirming the prognostic relevance of SEZ6L2 in colon cancer.

In conclusion, this comprehensive validation, spanning both mRNA and protein levels in a Chinese patient cohort, reaffirms the TCGA-based conclusion that SEZ6L2 is consistently overexpressed in colon cancer. The marked overexpression at the protein level emphasizes the potential functional relevance of SEZ6L2 in the context of colon cancer progression. These findings contribute not only to the understanding of SEZ6L2 as a molecular marker but also highlight the necessity of diverse patient cohorts for comprehensive cancer research. (Figure 2)

**Figure 2.**
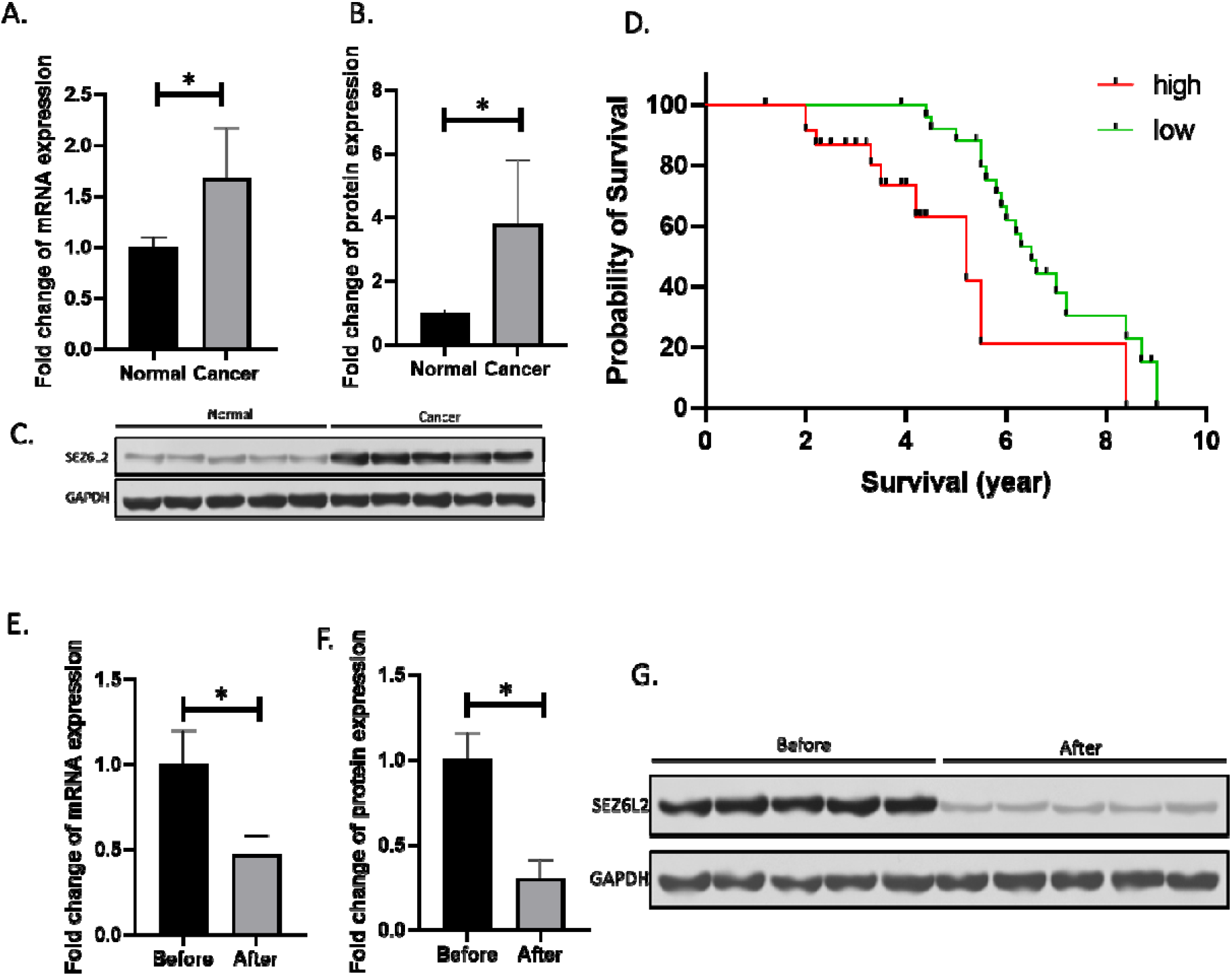
Validation of the survival association of SEZ6L2 in colon cancer and the effect of Bexarotene treatment. A. Fold change of mRNA expression of SEZ6L2 in colon cancer (n=77) and normal (n=49) colon tissues. B. Fold change of protein expression of SEZ6L2 in colon cancer (n=77) and normal (n=49) colon tissues. C. Representative images of western blotting for the protein expression of SEZ6L2 in colon cancer and normal colon tissues. D. Survival association of SEZ6L2 in colon cancer (n=50). E. Fold change of mRNA expression of SEZ6L2 in colon cancer before (n=50) and after (n=47) Bexarotene treatment (>90 days). F. Fold change of protein expression of SEZ6L2 in colon cancer before and after Bexarotene treatment (>90 days). G. Representative images of western blotting for the protein expression of SEZ6L2 in colon cancer before and after Bexarotene treatment (>90 days). *p<0.05

### 3.3. The effect of Bexarotene treatment on SEZ6L2

In our investigation into the pharmacological impact of Bexarotene on colon cancer, with a specific focus on its modulation of SEZ6L2 expression, we employed a cohort comprising patients both with and without Bexarotene treatment. Patients undergoing Bexarotene treatment for over 90 days were selected for analysis, while those without Bexarotene treatment, possibly undergoing alternative drug therapies, served as the control group. The study included 50 colon cancer patients before Bexarotene treatment and 47 patients after Bexarotene treatment, ensuring a comprehensive representation of the pharmacological effects of the drug.

Our analyses revealed a significant and consistent reduction in SEZ6L2 expression following Bexarotene treatment. Both mRNA and protein expression levels of SEZ6L2 exhibited a noteworthy decrease in patients subjected to Bexarotene compared to the control group without Bexarotene treatment. The average fold change in SEZ6L2 mRNA expression in colon cancer patients after Bexarotene treatment was significantly lower compared to those without treatment, highlighting the suppressive effect of Bexarotene on SEZ6L2 at the transcriptional level. Parallelly, large-scale protein expression analyses, conducted through Western blotting assays, further supported our findings. Patients treated with Bexarotene exhibited a substantial reduction in SEZ6L2 protein levels compared to those without Bexarotene treatment.

This comprehensive investigation provides compelling evidence that Bexarotene treatment is associated with a significant downregulation of SEZ6L2 in colon cancer patients. The observed reduction in both mRNA and protein levels underscores the pharmacological impact of Bexarotene on SEZ6L2 expression, suggesting its potential role as a targeted therapeutic approach in the management of colon cancer. These findings contribute valuable insights into the pharmacodynamics of Bexarotene and lay the groundwork for further exploration of SEZ6L2 as a promising pharmacological target in the context of colon cancer treatment. (Figure 2)

### 3.4. In vitro roles of SEZ6L2 in colon cancer cells

To delve deeper into the intricate interplay between SEZ6L2 and Bexarotene treatment, we conducted in vitro assays utilizing colon cancer cells. From the survival analysis, we observed that the survival impact of SEZ6L2 is evident across all four subtypes, with CMS3 displaying the most significant difference. Given that a KRAS mutation is a major characteristic of CMS3, employing cell lines with this mutation might yield the best results in in vitro studies. HCT116, which carries the G12D KRAS mutation[17], was therefore chosen for subsequent studies. Through the overexpression of SEZ6L2, we sought to elucidate its impact on both expression levels and functional aspects, specifically focusing on cell viability and migration.

Following the transfection of colon cancer cells with the SEZ6L2 plasmid, a concentration-dependent overexpression of SEZ6L2 was observed at both mRNA and protein levels. This successful modulation of SEZ6L2 expression provided a foundation for the subsequent exploration of its functional implications. The overexpression of SEZ6L2 was confirmed through meticulous assessments of mRNA and protein levels, establishing a direct correlation between plasmid concentration and SEZ6L2 expression.

Results from the CCK-8 assay revealed a noteworthy correlation between SEZ6L2 expression levels and cell viability. As SEZ6L2 expression increased in colon cancer cells, a concentration-dependent elevation in cell viability was observed. This suggests a potential role for SEZ6L2 in promoting cell viability. In contrast, despite the increase in SEZ6L2 expression, cell migration exhibited no significant differences. The modulation of SEZ6L2 did not exert a discernible impact on the migratory behavior of colon cancer cells.

Our in vitro assays shed light on the multifaceted role of SEZ6L2 in colon cancer. The concentration-dependent increase in SEZ6L2 expression correlated with enhanced cell viability, indicating a potential role in promoting cell survival. Notably, this effect was not mirrored in cell migration, suggesting that SEZ6L2 might play a more specific role in influencing cell viability rather than migratory behavior. These findings contribute to the nuanced understanding of the functional implications of SEZ6L2 in colon cancer cells. The observed effects on cell viability underscore the intricate interplay between SEZ6L2 expression levels and cellular functions, paving the way for further investigations into the underlying molecular mechanisms and potential therapeutic interventions targeting SEZ6L2 in the context of colon cancer. (Figure 3)

**Figure 3.**
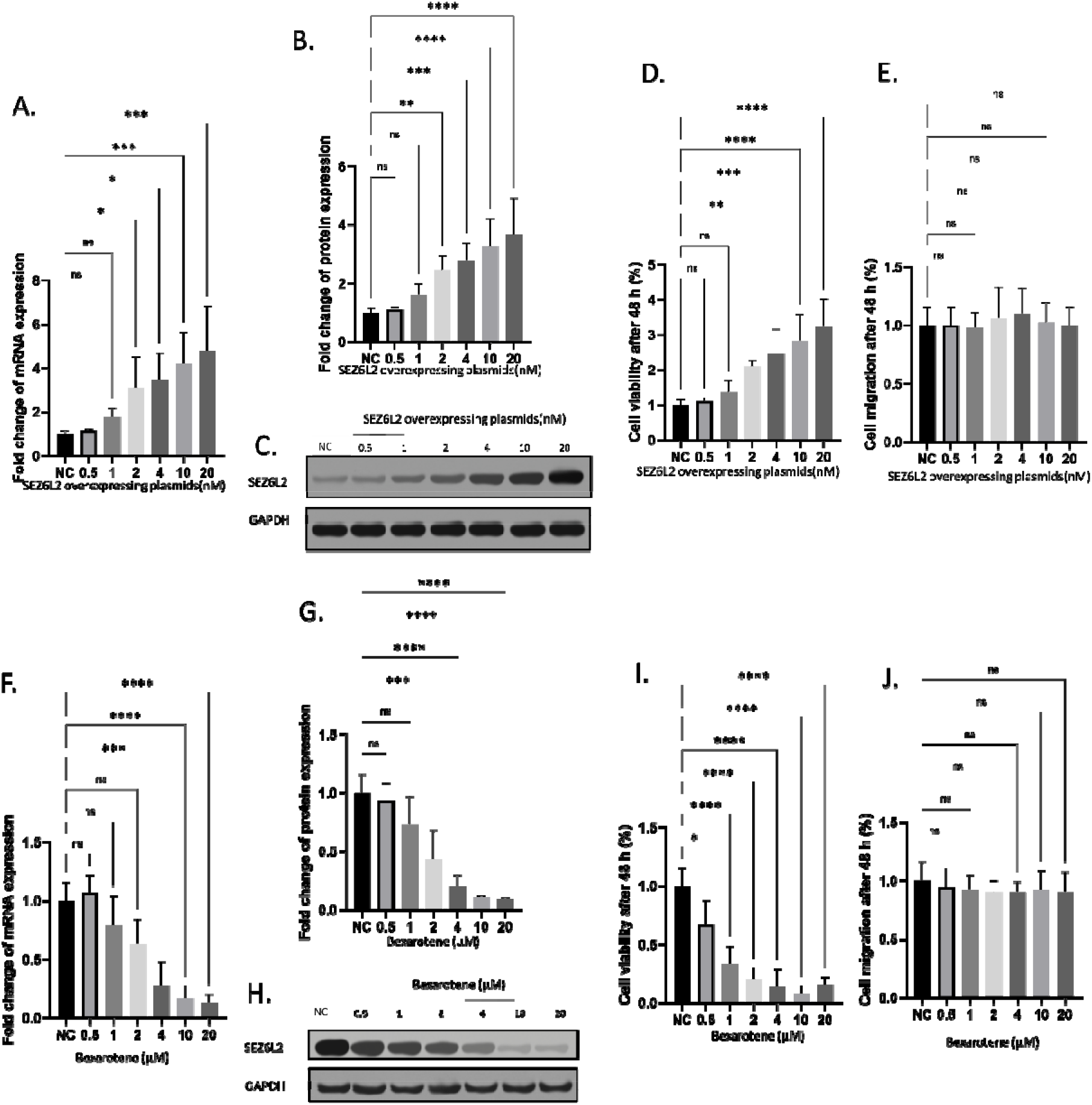
In vitro roles of SEZ6L2 in colon cancer cells HCT116 and Bexarotene treatment. All the data were normalized to the percentage of the control. A. Fold change of mRNA expression of SEZ6L2 in colon cancer cells with SEZ6L2 overexpression. B. Fold change of protein expression of SEZ6L2 in colon cance cells with SEZ6L2 overexpression. C. Representative images of western blotting fo the protein expression of SEZ6L2 in colon cancer cells with SEZ6L2 overexpression. D. Fold change of cell viability of colon cancer cells with SEZ6L2 overexpression. E. Fold change of cell migration of colon cancer cells with SEZ6L2 overexpression. F. Fold change of mRNA expression of SEZ6L2 in colon cancer cells with Bexarotene treatment. G. Fold change of protein expression of SEZ6L2 in colon cancer cells with Bexarotene treatment. H. Representative images of western blotting for the protein expression of SEZ6L2 in colon cancer cells with Bexarotene treatment. I. Fold change of cell viability of colon cancer cells with Bexarotene treatment. J. Fold change of cell migration of colon cancer cells with Bexarotene treatment.

### 3.5. In vitro effects of Bexarotene treatment on SEZ6L2 in colon cancer cells

Then, to investigate the in vitro effects of Bexarotene treatment on SEZ6L2 in HCT116 colon cancer cells, a range of concentrations (0.5 to 20 uM) was applied to assess its impact on SEZ6L2 expression, cell viability, and migration.

Results showed that Bexarotene exposure resulted in a dose-dependent reduction in both SEZ6L2 mRNA and protein expression in HCT116 cells. Notably, statistical significance was reached at higher doses, with SEZ6L2 mRNA expression significantly reduced when Bexarotene exceeded 4 uM, while protein expression showed significance at doses surpassing 2 uM.

Bexarotene treatment exhibited a dose-dependent decrease in colon cancer cell viability. Surprisingly, even at lower doses (0.5-2 uM) where SEZ6L2 expression did not show significant changes, HCT116 cell viability was already significantly lower than the control. This suggests that Bexarotene may influence cell viability through mechanisms beyond SEZ6L2 regulation. However, SEZ6L2 might be a major regulation for HCT116 cell viability. In contrast to its impact on viability, Bexarotene treatment, even at the highest concentration, did not elicit any significant effect on HCT116 cell migration. This indicates a specific influence on viability without concurrent alterations in migratory behavior.

These in vitro findings underscore the dose-dependent inhibitory effect of Bexarotene on SEZ6L2 expression in HCT116 cells at both mRNA and protein levels. The observed reduction in HCT116 cell viability, even at lower Bexarotene doses where SEZ6L2 expression remained unchanged, suggests additional mechanisms contributing to the drug’s impact on cell survival. Notably, the lack of influence on cell migration highlights the specificity of Bexarotene’s effects on distinct cellular processes. (Figure 3)

### 3.6. Validation of In vitro roles of SEZ6L2 in colon cancer cells using HCT15

To alidate the In vitro roles of SEZ6L2 in colon cancer cells, we investigated the in vitro effects of Bexarotene treatment on SEZ6L2 in HCT15 colon cancer cells, in a parallel experiments as we did for HCT116.

Results showed a very similar results in HCT15 as in HCT116 that Bexarotene exposure resulted in a dose-dependent reduction in both SEZ6L2 mRNA and protein expression in colon cancer cells. Higher doses of Bexarotene were found to significantly reduce SEZ6L2 mRNA levels when the concentration reached over 4 uM, and protein levels were similarly affected at doses over 2 uM. Additionally, Bexarotene demonstrated a dose-responsive reduction in the viability of colon cancer cells. Interestingly, even at lower concentrations (0.5-2 uM), where there was no significant change in SEZ6L2 expression, there was a notable decrease in HCT15 cell viability compared to the control group. This implies that Bexarotene’s effect on cell survival may operate through pathways other than just SEZ6L2 regulation. Nonetheless, SEZ6L2 could be a critical factor in the viability of HCT15 cells. Despite its significant impact on cell viability, Bexarotene did not significantly affect colon cancer cell migration at any concentration tested, indicating that its effects are specific to HCT15 cell survival rather than affecting HCT15 cell migratory capabilities.

These laboratory results highlight Bexarotene’s dose-dependent suppressive effects on SEZ6L2 expression at both the mRNA and protein levels within HCT15 cells. The reduction in viability observed at lower doses, without changes in SEZ6L2 expression, points to other mechanisms at play that influence the drug’s effect on HCT15 cell survival. The absence of effects on cell migration emphasizes Bexarotene’s targeted action on specific cell functions. These findings offer important insights into the complex relationship between Bexarotene, SEZ6L2, and cell functions in colon cancer, laying the groundwork for further research into the molecular pathways involved and its potential therapeutic uses in treating colon cancer. (Figure 4)

**Figure 4.**
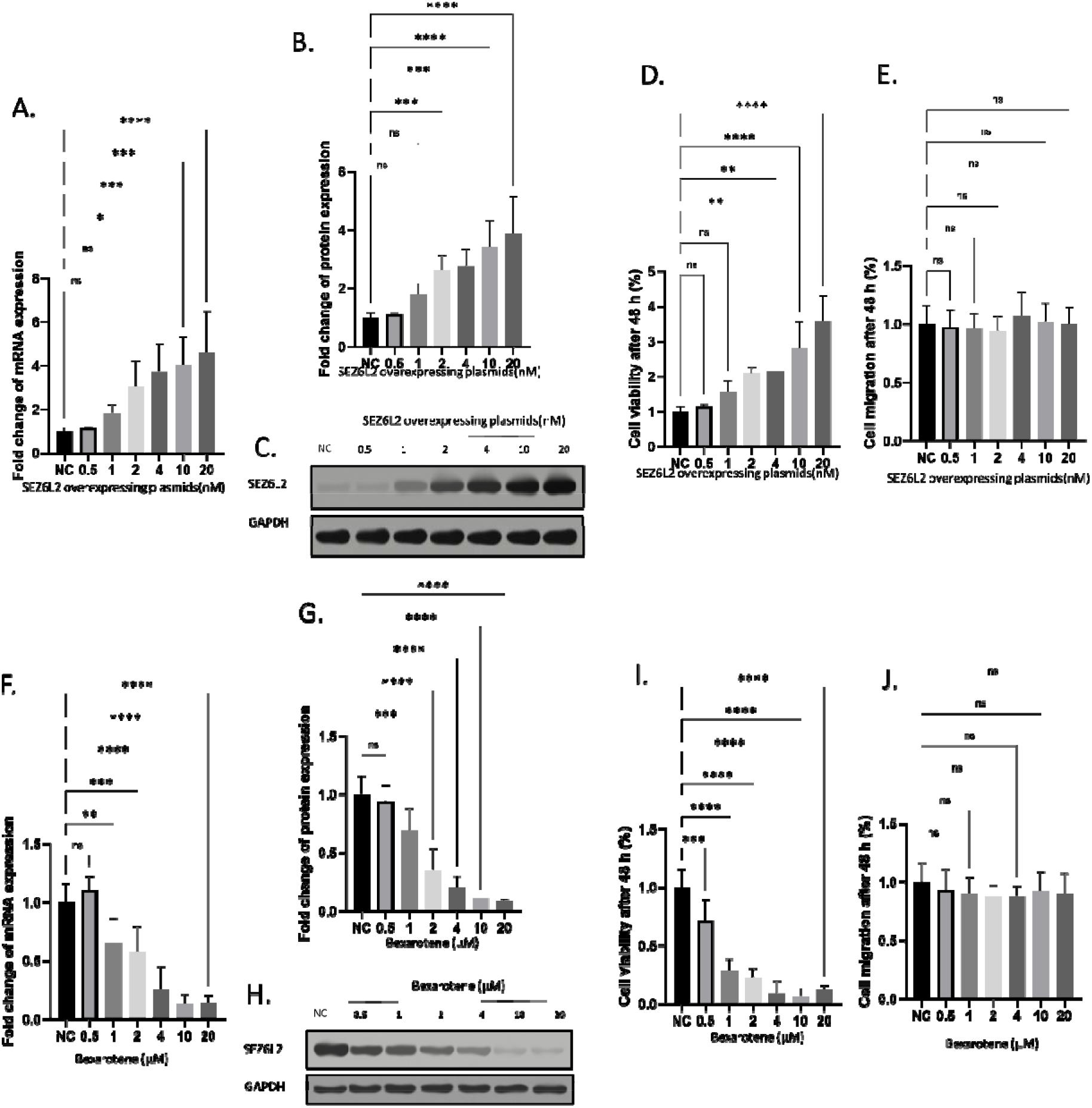
Validation of In vitro roles of SEZ6L2 in colon cancer cells using HCT15. All the data were normalized to the percentage of the control. A. Fold change of mRNA expression of SEZ6L2 in colon cancer cells with SEZ6L2 overexpression. B. Fold change of protein expression of SEZ6L2 in colon cancer cells with SEZ6L2 overexpression. C. Representative images of western blotting for the protein expression of SEZ6L2 in colon cancer cells with SEZ6L2 overexpression. D. Fold change of cell viability of colon cancer cells with SEZ6L2 overexpression. E. Fold change of cell migration of colon cancer cells with SEZ6L2 overexpression. F. Fold change of mRNA expression of SEZ6L2 in colon cancer cells with Bexarotene treatment. G. Fold change of protein expression of SEZ6L2 in colon cancer cells with Bexarotene treatment. H. Representative images of western blotting for the protein expression of SEZ6L2 in colon cancer cells with Bexarotene treatment. I. Fold change of cell viability of colon cancer cells with Bexarotene treatment. J. Fold change of cell migration of colon cancer cells with Bexarotene treatment.

### 3.7. In vivo roles of SEZ6L2 in colon cancer

In our pursuit to unravel the impact of SEZ6L2 on colon cancer progression, we extended our investigation to an in vivo setting by employing a colon cancer animal model. To establish this model, colon cancer cells were transfected with a construct incorporating SEZ6L2 and a G418 resistance gene. Following transfection, cells were subjected to G418 selection to generate a stable cell line expressing SEZ6L2. The parental cell line, devoid of SEZ6L2 overexpression, served as the control in the animal model.

Stable cell lines expressing SEZ6L2 were successfully generated through G418 selection, ensuring the sustained overexpression of SEZ6L2 in the experimental group. The parental cell line, used as a control, lacked this overexpression. Animals were subjected to the respective colon cancer cell lines to observe the in vivo consequences of SEZ6L2 overexpression. Survival data were meticulously recorded to discern any potential disparities in the survival rates between the experimental group (SEZ6L2-overexpressing cells) and the control group (parental cells).

Notably, our findings revealed a significant difference in the survival rates of animals modeled with colon cancer cells overexpressing SEZ6L2 compared to those with the control cells. Animals in the experimental group exhibited a markedly accelerated decline in survival, succumbing to the disease at a much faster rate than their counterparts in the control group.

This in vivo exploration underscores the crucial role of SEZ6L2 in influencing the aggressiveness of colon cancer. The accelerated decline in survival rates observed in animals modeled with SEZ6L2-overexpressing cells emphasizes the potential of SEZ6L2 as a prognostic indicator and therapeutic target in colon cancer. These results provide valuable insights into the in vivo implications of SEZ6L2 overexpression and warrant further investigation into the underlying molecular mechanisms driving its impact on colon cancer progression. (Figure 5)

**Figure 5.**
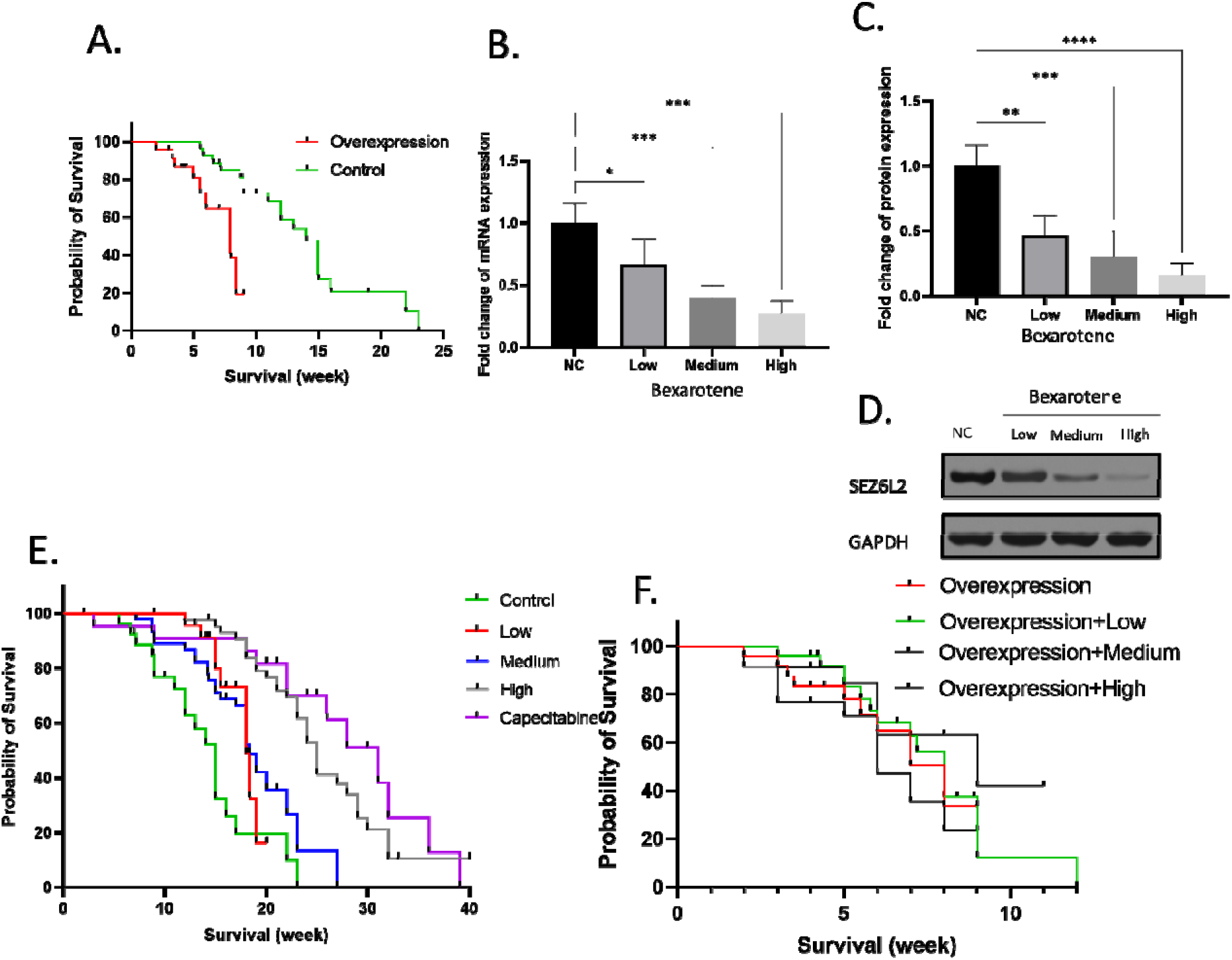
In vivo roles of SEZ6L2 in colon cancer and Bexarotene treatment. A. Effect of SEZ6L2 overexpression on animal survival. B. Fold change of mRNA expression of SEZ6L2 in colon cancer tissue from animals with Bexarotene treatment. C. Fold change of protein expression of SEZ6L2 in colon cancer tissue from animals with Bexarotene treatment. D. Representative images of western blotting for the protein expression of SEZ6L2 in colon cancer tissue from animals with Bexarotene treatment. E. Effect of Bexarotene treatment on animal survival. F. Effect of Bexarotene treatment on SEZ6L2 overexpressing animal survival.

### 3.8. In vivo effects of Bexarotene treatment on SEZ6L2 in colon cancer

Next, in our ongoing efforts to establish the therapeutic impact of Bexarotene on colon cancer, we employed an in vivo approach using a colon cancer animal model. For this experiment, we utilized the parental colon cancer cell line for consistency and treated the animals with varying doses of Bexarotene (Low, Medium, and High).

Our results demonstrated a significant and dose-dependent reduction in the expression of SEZ6L2 at both the mRNA and protein levels following Bexarotene treatment. All three doses of Bexarotene treatment (Low, Medium, and High) led to a significant reduction in SEZ6L2 mRNA levels in comparison to the untreated control group. Correspondingly, Bexarotene treatment resulted in a decrease in SEZ6L2 protein expression. The reduction was evident across all three doses, reinforcing the consistency of the observed effect.

Our in vivo experiments provide compelling evidence that Bexarotene treatment exerts a robust suppressive effect on SEZ6L2 expression in colon cancer. The significant reduction observed in both mRNA and protein levels following Bexarotene administration underscores its potential as a therapeutic agent for modulating SEZ6L2 expression in the context of colon cancer. These findings contribute valuable insights into the molecular responses to Bexarotene treatment and support its potential utility as a targeted therapeutic approach for managing colon cancer by modulating SEZ6L2 expression. Further investigations are warranted to elucidate the detailed mechanisms underlying this observed effect and to evaluate the broader implications of Bexarotene in colon cancer therapy. (Figure 5)

### 3.9. In vivo effects of SEZ6L2 on Bexarotene treatment toward colon cancer

In addition, to comprehensively evaluate the therapeutic efficacy of Bexarotene in colon cancer, we conducted animal experiments employing two distinct experimental setups. In the control treatment experiment, parental colon cancer cells were utilized, while in the second experiment, animals were modeled with stable SEZ6L2-overexpressing colon cancer cells.

In the control treatment experiment, animals modeled with parental colon cancer cells were treated with three different doses of Bexarotene. The results demonstrated a notable increase in the survival rate of colon cancer animals with Bexarotene treatment. Although the differences between the various dose groups were not starkly pronounced, the overall trend indicated a positive impact on survival.

Contrastingly, in the experimental setup involving animals modeled with stable SEZ6L2-overexpressing colon cancer cells, Bexarotene treatment failed to elicit a discernible difference in animal survival. The survival curves for Bexarotene High (H), Medium (M), Low (L) groups closely mirrored that of the control group (no drug), indicating a lack of therapeutic efficacy in this specific context. In addition, we applied first-line drug Capecitabine as the positive control, result showed that the high Bexarotene achieved similar results as the Capecitabine.

Our findings underscore the context-dependent nature of Bexarotene’s therapeutic effects in colon cancer. While Bexarotene demonstrated a positive impact on the survival of animals modeled with parental colon cancer cells, this effect was not recapitulated in the SEZ6L2-overexpressing model. The close alignment of survival curves in the SEZ6L2 overexpressing model, regardless of Bexarotene treatment, suggests a potential interaction or modulation of Bexarotene’s effects by SEZ6L2 overexpression. These results highlight the importance of considering specific molecular contexts in assessing the efficacy of therapeutic interventions, emphasizing the need for further investigations to elucidate the intricate interplay between Bexarotene, SEZ6L2, and colon cancer progression. (Figure 5)

### 3.10. Direct interaction between SEZ6L2 protein and Bexarotene

To explore the putative direct interaction between SEZ6L2 protein and Bexarotene, we employed computational prediction methods leveraging the chemical structure of Bexarotene and the predicted structure of SEZ6L2 protein obtained from the AlphaFold database. The confidence in the AlphaFold model was substantiated by a prediction error plot, highlighting high confidence, particularly in the central region of the protein—our region of interest for potential Bexarotene binding.

Utilizing the obtained chemical structure of Bexarotene and the predicted structure of SEZ6L2, we conducted protein-ligand binding modeling to predict the potential interaction. The results revealed a model with a relatively high affinity, as indicated by a calculated energy score lower than −8.3 kJ/mole.

Our computational predictions strongly suggest the existence of a direct interaction between SEZ6L2 protein and Bexarotene. The high confidence in the AlphaFold model, coupled with the favorable energy score obtained from the protein-ligand binding modeling, points towards a potential and relatively high-affinity binding between SEZ6L2 and Bexarotene. These computational insights provide a basis for further experimental validation and mechanistic exploration of the direct interaction between SEZ6L2 and Bexarotene. The predicted model offers a starting point for understanding the molecular basis of their interaction and may guide future studies on the development of targeted therapeutic strategies in the context of colon cancer treatment (Figure 6).

**Figure 6.**
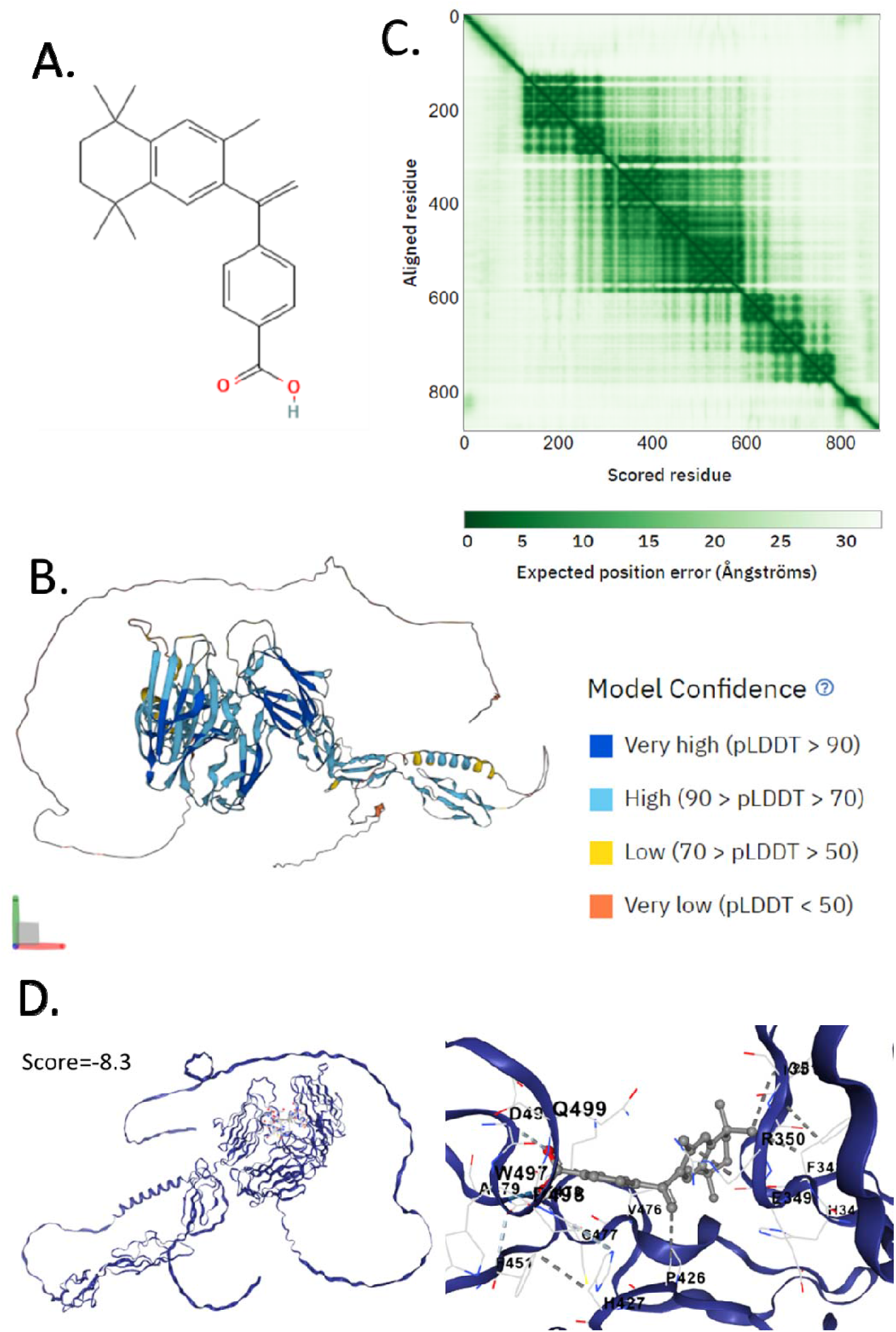
Direct interaction between SEZ6L2 protein and Bexarotene. A. Chemical structure of Bexarotene. B. Alpha-fold predicted structure of SEZ6L2 protein. C. Alpha-fold model position error. D. binding model between SEZ6L2 protein and Bexarotene.

## 4. Discussion

Colon cancer remains a major global health concern, necessitating continuous efforts to unravel its molecular intricacies and develop effective therapeutic strategies. SEZ6L2 has been reported to promote the growth and metastasis of breast cancer[18]. In this study, we focused on SEZ6L2 as a potential biomarker and therapeutic target for colon cancer. Our comprehensive analyses, spanning clinical databases, in vitro experiments, and in vivo animal models, collectively contribute valuable insights into the multifaceted role of SEZ6L2 in colon cancer.

In contrast to other investigations in the field, the exploration of medical drugs and bioinformatics has advanced our understanding of the mechanisms and treatment strategies for numerous human diseases [19–32], especially for drugs that might have multiple effects[22,25,27,33–36]. The identification of targets of drugs is one of the challenges in pharmacological studies[37]. Genetic biomarker discovery from TCGA has been previous conducted[30,38–46]. Our initial analysis of TCGA and GTEx databases revealed a consistent and robust overexpression of SEZ6L2 in colon cancer tissues compared to normal colon tissues. This overexpression was validated in a cohort of Chinese patients, emphasizing the importance of considering ethnic diversity in cancer research. The previous study using TCGA to study the prognostic role of this gene focused on white patients[7,28,29,47] that are different from Chinese patients in our study. The diagnostic potential of SEZ6L2 was underscored by a high AUC in ROC analysis, suggesting its viability as a diagnostic biomarker for colon cancer. Furthermore, survival analysis indicated a significant correlation between high SEZ6L2 expression and lower survival rates, positioning SEZ6L2 as a promising prognostic biomarker. These findings collectively highlight the clinical relevance of SEZ6L2 in the context of colon cancer diagnosis and prognosis. SEZ6L2 has been previously suggested to be useful for thyroid carcinoma prognosis[48]. Overexpression of SEZ6L2 also found to predict poor prognosis in patients with cholangiocarcinoma[49]. Simimar prognostic biomarker result is also found in lung cancer[50]. All these previous studies were consistent with our conclusion that it might be a practical prognosis biomarker for clinical applications.

The effectiveness of the Bexarotene in reducing drug resistance has been observed through its regulation of RFX1 in embryonic carcinoma cells[51]. Applying physiologically based pharmacokinetic modeling, a previous study extrapolate the metabolism of bexarotene in the context of chronic kidney disease and acute kidney injury in rats[52]. However, the specific mechanisms of Bexarotene in relation to colon cancer are not well-defined, especially concerning whether Bexarotene directly influences colon cancer cells. To explore the potential therapeutic implications of SEZ6L2, we investigated its response to Bexarotene treatment. Our analyses revealed a significant downregulation of SEZ6L2 at both mRNA and protein levels in colon cancer patients following Bexarotene treatment. This pharmacological impact was further validated in in vitro experiments, where Bexarotene treatment led to a dose-dependent reduction in SEZ6L2 expression and cell viability. Interestingly, the inhibitory effect on cell viability was observed even at lower Bexarotene doses where SEZ6L2 expression remained unchanged, suggesting additional mechanisms contributing to the drug impact on cell survival. These findings provide a foundation for considering SEZ6L2 as a potential pharmacological target in colon cancer treatment.

As showen in the western blooting, the colon cancer cell line expressed low level of SEZ6L2 and we believe that the knockout might not achieve the best results. Hence, we only conducted overexpression experiments. In our in vivo experiments, SEZ6L2 overexpression in colon cancer cells significantly accelerated the decline in animal survival rates. This observation underscores the potential of SEZ6L2 as a prognostic indicator for colon cancer aggressiveness. Additionally, Bexarotene treatment exhibited a dose-dependent reduction in SEZ6L2 expression in vivo, supporting its potential as a therapeutic agent for modulating SEZ6L2 levels in colon cancer. However, the therapeutic efficacy of Bexarotene was influenced by SEZ6L2 overexpression, as animals with SEZ6L2-overexpressing cells did not exhibit a significant improvement in survival with Bexarotene treatment. This highlights the need for a more nuanced understanding of the interplay between SEZ6L2 expression and therapeutic responses to optimize treatment strategies.

Our in vitro assays exploring the functional implications of SEZ6L2 in colon cancer cells revealed a concentration-dependent increase in cell viability with SEZ6L2 overexpression. However, this effect was not mirrored in cell migration, suggesting a more specific role for SEZ6L2 in influencing cell viability rather than migratory behavior. These findings contribute to the nuanced understanding of the functional implications of SEZ6L2 in colon cancer cells, paving the way for further investigations into the underlying molecular mechanisms. Previous findings revealed that silencing SEZ6L2 significantly curtailed breast cancer cell proliferation and triggered cell cycle arrest in the G1 phase[53]. Moreover, the suppression of SEZ6L2 reduced cell migration and invasion. Conversely, overexpressing SEZ6L2 had the reverse effects[53]. SEZ6L2 also enhanced tumorigenesis in breast cancer cells in vivo. Additionally, using bioinformatics tools, upstream transcription factor 1 (USF1) was identified as a regulator that binds to the SEZ6L2 promoter and positively influences its transcription[53]. Overall, this study demonstrated that SEZ6L2, under the regulatory control of USF1, plays a crucial role in the proliferation and metastasis of breast cancer cells[53]. Understanding SEZ6L2 function in BC adds to our knowledge of its pathogenesis and may provide insights beneficial for developing breast cancer therapies[53]. Our data suggested that, this gene play roles in migration not only in breast cancer cells but also in colon cancer cells.

While our study provides valuable insights into the diagnostic, prognostic, and therapeutic potential of SEZ6L2 in colon cancer, several questions and avenues for future research emerge. A previous study suggested that Bexarotene-induced cell death in ovarian cancer cells is through Caspase-4-gasdermin E mediated pyroptosis [54]. Another study suggested that the optimized Bexarotene Aerosol formulation inhibits major subtypes of lung cancer in mice[55]. These study provide background for the current study. The precise mechanistic involvement of SEZ6L2 in colon cancer progression remains to be elucidated. Further exploration of its molecular interactions, downstream signaling pathways, and potential crosstalk with other molecules will enhance our understanding of its role in the complex landscape of colon cancer. Additionally, the influence of SEZ6L2 on the therapeutic response to Bexarotene warrants detailed investigation, with a focus on optimizing treatment strategies for individuals with varying SEZ6L2 expression levels. The potential use of SEZ6L2 as a therapeutic target in colon cancer may open avenues for the development of novel targeted therapies with enhanced efficacy and specificity. In addition, diseases of the digestive system might be associated with microbiota[27,35,36]. Whether microbiota are associated with SEZ6L2 remains unknown and requires further exploration. Understanding this potential association could shed light on the role of microbiota in digestive system diseases and their impact on SEZ6L2, potentially opening new avenues for research and treatment strategies.

In conclusion, our study positions SEZ6L2 as a compelling biomarker for colon cancer diagnosis and prognosis, with potential therapeutic implications. The integration of clinical, in vitro, and in vivo data strengthens the translational relevance of our findings, offering a comprehensive view of SEZ6L2 in the context of colon cancer. As we continue to unravel the molecular intricacies of colon cancer, the insights gained from this study contribute to the ongoing efforts to improve patient outcomes through personalized and targeted therapeutic approaches.

## Author Contributions

Huajun Zheng and Jianying Zheng contributed equally to this work. Huajun Zheng and Jianying Zheng conceptualized the project; Huajun Zheng was responsible for the material preparation and characterization; Huajun Zheng, Jianying Zheng, and Yan Shen conducted the in vitro experiments; Jianying Zheng performed experiments in vivo; Yan Shen performed bioinformatic analysis and provided guidance on experimental design and wrote the manuscript.

## Competing interest

The authors declare no competing financial interest.

## Acknowledgments

This work is supported by The Second Affiliated Hospital of Zhejiang University of Traditional Chinese Medicine.

## Notes

### Competing Interest Statement

The authors have declared no competing interest.

